# A biological source of marine sedimentary iron oxides

**DOI:** 10.1101/108621

**Authors:** Jacob P. Beam, Jarrod J. Scott, Sean M. McAllister, Clara S. Chan, James McManus, Filip J. R. Meysman, David Emerson

## Abstract

The biogeochemical cycle of iron is intricately linked to numerous element cycles. Although reductive biological processes that bridge the iron cycle to other element cycles are established, little is known about microbial oxidative processes on iron cycling in sedimentary environments—resulting in the formation of iron oxides. Here, we show that a major source of sedimentary iron oxides originates from the metabolic activity of iron-oxidizing bacteria from the class Zetaproteobacteria, stimulated by burrowing animals in coastal sediments. Zetaproteobacteria were estimated to be a global total of 10^26^ cells in coastal, bioturbated sediments and would equate to an annual production of approximately 7.9 x 10^15^ grams of sedimentary iron oxides—twenty-five times larger than the annual flux of iron oxides by rivers. These data suggest that iron-oxidizing Zetaproteobacteria are keystone organisms in marine sedimentary environments given their low numerical abundance; yet exert a profound impact via the production of iron oxides.

## Main

Iron oxides are important components of coastal and continental shelf sediments, and are thought to originate primarily by river deposition (Poulton and Raiswell, 2002). Authigenic iron oxides can also be formed by the oxidation of ferrous iron [Fe(II)], which has been largely attributed to chemical oxidation of Fe(II) in pore waters in areas with significant sediment mixing and irrigation by animals—bioturbation and bioirrigation—and subsequent reaction with oxygen in anoxic sediments (Aller, 1982; Canfield, 1989). Although sedimentary chemical oxidation of iron is important under saturated oxygen conditions at neutral pH (Canfield, 1989), bioirrigated sediments contain microenvironments—formed by burrowing animals—that are well below saturation (Kristensen and Kostka, 2005) where the biological contribution to iron oxidation is quantitatively more significant (Emerson *et al.*, 2010). Under low-oxygen conditions (< 100 μM O_2_) and without rapid replenishment of highly reactive, poorly-crystalline iron oxides (i.e., ferrihydrite and lepidocrocite), they would be quickly exhausted by hydrogen sulfide and by bacterial iron reduction, and form iron sulfides or be released to the water column. Collectively, these findings suggest that biology is involved and important in marine sedimentary iron oxidation under ferruginous conditions commonly observed in bioturbated sediments.

The Zetaproteobacteria represent a class of iron-oxidizing bacteria (FeOB) that are exclusively found in marine or saline-influenced environments that contain high ferrous iron [Fe(II)] concentrations (McAllister *et al.*, 2011; McBeth *et al.*, 2013; Scott *et al.*, 2015, 2017). Coastal marine sediments can have Fe(II) pore water concentrations ranging from ∼1-2,000 μmol L^-1^ that are capable of supporting lithoautotrophic populations of Zetaproteobacteria (Emerson, 2016). Recent studies have identified Zetaproteobacteria in surface openings of benthic macrofauna in the Mediterranean Sea (Rubin-Blum *et al.*, 2014), worm burrows in sub-marine groundwater discharge into sands in Delaware (McAllister *et al.*, 2015), coastal sediments in Denmark (Laufer *et al.*, 2016), and Baltic and North Sea sediments (Reyes *et al.*, 2016). These recent studies suggest that Zetaproteobacteria may play a significant role in iron oxidation in marine sediments—a quantitative estimate of their abundance is necessary to determine their biogeochemical role on a global scale.

We analyzed the coastal sediment microbial communities to determine the extent of Zetaproteobacteria from geographically diverse sites (n=90; Supplementary Table 1) utilizing 16S rRNA gene sequencing, which highlights their importance on a global scale. Iron oxidation appears ubiquitous in the Zetaproteobacteria (Field *et al.*, 2015; Scott *et al.*, 2015; Barco *et al.*, 2015), thus the 16S rRNA gene can be used to infer this specific metabolism. We also enriched for environmentally relevant iron-oxidizing bacteria from coastal sediments, which provided further metabolic evidence of the importance of iron oxidation in marine sediments. A meta-analysis of 16S rRNA gene studies revealed the extent and importance of Zetaproteobacteria on the global sedimentary iron biogeochemical cycle.

We identified Zetaproteobacteria in 60% of our samples (Supplementary Table S1), and their median relative abundance was 1.1 percent of the total microbial community (range = 0.04-15%) in worm (e.g., polychaetes) burrows in coastal marine sediments (Fig. 1a). Zetaproteobacteria were ten times less abundant in bulk sediments (Fig. 1a) with a median near zero percent (range = 0-1%), and were statistically different from worm burrows (p-value=9.2 x 10-7, Wilcoxon test). The large ranges (0.04-15%) and non-normal distribution of Zetaproteobacteria relative abundance in worm burrows (see Supplementary Materials and Methods) was most likely a combination of differences in burrow ventilation rates and efficiencies (Kristensen and Kostka, 2005), differences in sediment physicochemical conditions (Supplementary Table S1), and sampling bias (for example, residual sediment on worm burrows). Bioirrigation by benthic animals increases the extent of oxidative processes in these sediments, thus biological iron oxidation can occur at greater depths (10s of centimeters) than typical oxygen penetration of a few millimeters into coastal surface sediments. The abundance of Zetaproteobacteria at the burrow walls correlated with the concentration of pore water ferrous iron [Fe(II)] (Fig. 1b), which is their main energy source—resulting in the production of solid phase iron oxides around worm burrows (Fig. 1c). Quantitatively, highly reactive, iron oxides— operationally extractable by sodium dithionite (Poulton and Canfield, 2005)—were 3 times higher at burrow walls, which accounted for 20-40% of the iron oxides with depth (Supplementary Figure S1 and S2). These freshly-produced iron oxides are important substrates for iron-reducing microorganisms that release Fe(II) into pore waters, and are essential to the supply of dissolved iron (dFe) to phytoplankton in the water column in coastal communities and continental shelves (Severmann *et al.*, 2010). Iron oxides are also important to the mineralization of organic matter in marine sediments by iron reducers (Canfield, 1989) substrates for early pyrite diagenesis (Berner, 1984), enhance organic matter burial (Lalonde *et al.*, 2012), and inhibit accumulation of pore water hydrogen sulfide, preventing conditions detrimental to benthic animals (Kristensen and Kostka, 2005), perhaps functioning as a local firewall against euxinic conditions(Seitaj *et al.*, 2015).

**Figure 1.**
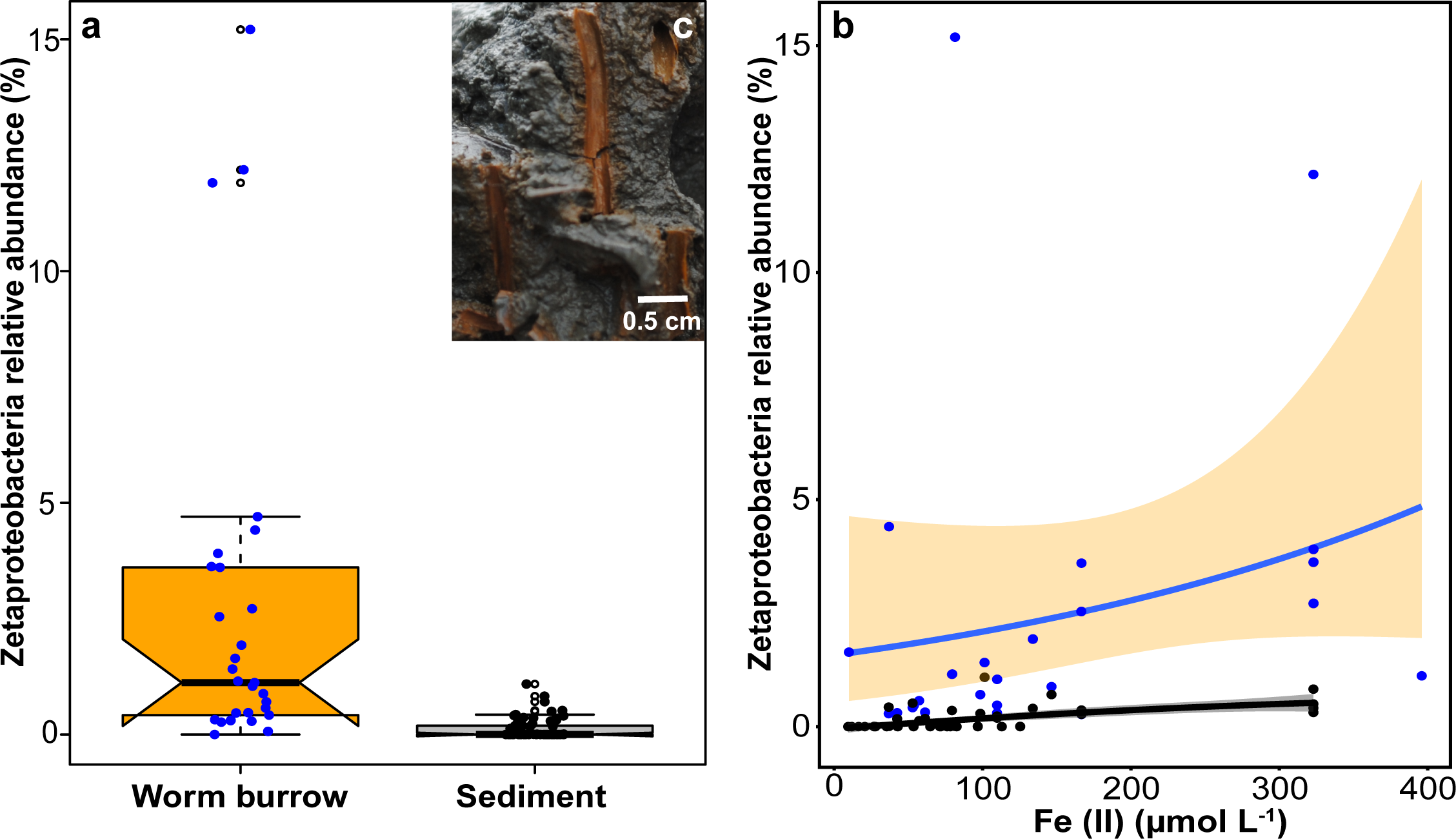
Boxplots of the relative abundance of Zetaproteobacteria (**a**) in iron-oxide lined worm burrow walls (n=29) and surrounding sediments (n=61). Notches are representative of 95% confidence interval and the medians (solid black lines) between worm burrows and sediments (1.1% and 0%, respectively) are statistically different (p-value=9.2 x 10^−7^, Wilcoxon test). Filled circles represent individual data points and open circles indicate outliers. Zetaproteobacteria relative abundance (‥) as a function of pore water ferrous iron [Fe(II)] concentration (μmol L-1) (**b**) from worm burrows (blue circles, fitted blue line, orange fill = 95% confidence interval) and sediments (black circles, fitted black line, grey fill = 95% confidence interval) (see Methods for details on line fits). Characteristic iron oxide lined worm burrow walls (c) from “The Eddy”, Sheepscot River, Maine, USA (image from 27 August 2015). Burrow walls are likely created by the polychaete, *Nereis diversicolor* or hemichordate, *Saccoglossus kowalevskii*, which are both common to these intertidal sediments in Maine.

The relative abundance of Zetaproteobacteria in worm burrows resembles the Fe(II) concentration profile (Supplementary Figure S3), and both were at their maximum values around 2-3 cm. The high Fe(II) (∼40-140 μM) and low oxygen (∼20-60 μM) conditions present in bioturbated sediment pore waters (Supplementary Figure S3) were ideal habitats for microaerophilic Zetaproteobacteria to thrive (Emerson *et al.*, 2010). Zetaproteobacteria relative abundance decreased with depth in both burrows and sediments (Supplementary Fig. S3), likely due to the decrease in Fe(II) with depth and increase in hydrogen sulfide production by sulfate-reducing bacteria. Although there is oxygen in these sediments at depth, hydrogen sulfide may inhibit oxygen respiratory machinery under these conditions. The formation of iron sulfide minerals with increasing depth by biogenic sulfide may also compete with Zetaproteobacteria for access to Fe(II). Under these ferruginous settings, biotic rates of Fe(II) oxidation exceed abiotic chemical oxidation (Emerson *et al.*, 2010).

Two Zetaproteobacteria Operational Taxonomic Units (herein, referred to as ZetaOTUs) dominated the Zetaproteobacterial diversity in worm burrows (Supplementary Table S2). The dominant ZetaOTU across all samples was ZetaOTU14, which comprised 32% of all ZetaOTUs (Supplementary Table S2), and is represented by four single cell amplified genomes (SAGs) from diffuse flow vent systems (Field *et al.*, 2015; Scott *et al.*, 2015, 2017). We isolated the first member of ZetaOTU14, strain CSS-1 from iron oxide surface flocculent in a laboratory bioturbation microcosm (Supplementary Figure S4). This strain grew best under low oxygen (∼60 μM O_2_) and high Fe(II) concentrations similar to those measured from sediment pore waters (Supplementary Figure S3). Strain CSS-1 produced stalks encrusted with poorly-crystalline iron oxides under laboratory conditions (Supplementary Figure S4). These iron oxides are consistent with those produced by other Zetaproteobacteria (Chan *et al.*, 2010), as well as in naturally occurring iron mats associated with hydrothermal vents, which are highly reactive, and resistant to undergoing diagenesis to more crystalline oxides (e.g., goethite) (Picard *et al.*, 2015). Single cell genomes from ZetaOTU14 representatives contained genes essential for growth on iron and low oxygen conditions (Supplementary Table S2). The second most abundant OTU was ZetaOTU9 (22%) and is represented by two cultured isolates (*Ghiorsea bivora* strains TAG-1 and SV108) (Mori *et al*.,, 2017), as well as 5 SAGs from deep-sea vents (Supplementary Table S2). ZetaOTU9 isolates also had genes necessary for growth on iron and low oxygen (Supplementary Table S2), and have also been shown to oxidize hydrogen, which may explain the ubiquity of this OTU in sediments and other environments (see below). There was no clear distribution of ZetaOTUs 14 and 9 with respect to depth (Supplementary Figure S3) in worm burrows and sediments—although it is likely that they are adapted for specific Fe(II) and O_2_ concentrations, which were hypothesized for other Zetaproteobacteria (Field *et al.*, 2015).

We searched for Zetaproteobacterial 16S rRNA gene sequences in marine sediment datasets (Supplementary Table S3), and identified them in numerous sediments on a global scale (Figure 2). We found a pattern consistent with our samples—ZetaOTUs 14 and 9 were present and generally the most abundant ZetaOTUs in coastal and shelf sediments (Figure 2). Zetaproteobacteria relative abundance was not found to exceed one percent in other studies, as microenvironments were not considered, which are abundant in bioturbated sediments (Kristensen and Kostka, 2005). Accordingly, we hypothesize that when the abundance of Zetaproteobacteria exceeds ∼0.1% in sediments, there is active growth and iron oxidation associated with bioturbating and bioirrigating animals. We estimated a median global population size of Zetaproteobacteria to be 1.05 x 10^26^ cells (range = 3.83 x 10^24^-1.44 x 10^27^ cells) from our measurements (Fig. 1a) and from other studies (Fig. 2) in continental shelf sediments using cellular abundance in the upper 10 centimeters—the worldwide average depth of bioturbation (Boudreau, 1998)—of continental shelf environments (total cells =10^29^; <150 m water depth (Kallmeyer *et al.*, 2012)). The global Zetaproteobacteria abundance estimate was then used with recent iron oxide production rate measurements from diffuse flow vents (∼1.3 x 10^−16^ mol Fe cell^-1^ hr^-1^) (Emerson *et al.*, 2016), which could result in the production of ∼7 petagrams of iron oxides per year (range = 0.1-70 petagrams per year). Recent two-dimensional, sub-millimeter Fe(II) measurements in bioturbated sediments revealed extensive Fe(II) oxidation occurring within the immediate vicinity of worm burrows and a rapid re-oxidation rate of 3.78 ±1.4 mmol Fe m^-2^ day^-1^ (de Chanvalon *et al.*, 2017). These chemical rate measurements combined with an estimate of the global volume of bioturbated coastal sediments 10 cm deep (∼2.1 x 10^13^ m3) (Teal *et al.*, 2008) would equate to an annual production of 1.6 ± 1.1 petagrams of iron oxides. These two independent estimations of iron oxide production rates are well within the range of one another. Based on these estimates, the annual biological oxidation of iron in sediments— forming iron oxides—could exceed the annual flux of iron oxides from rivers to coastal sediments (Poulton and Raiswell, 2002) up to a factor of twenty-five.

**Figure 2.**
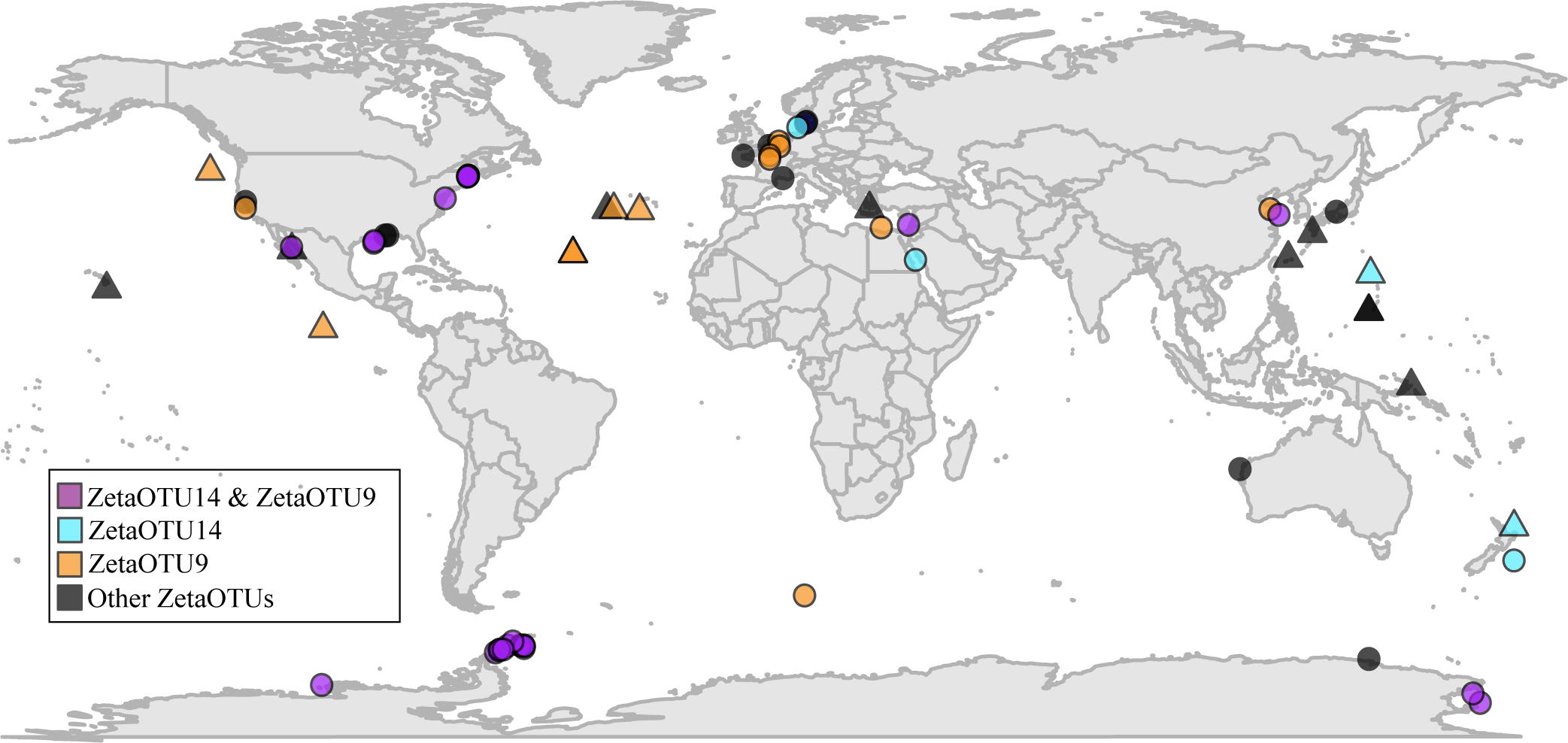
Global distribution of Zetaproteobacteria in marine sediments (circles) and non-sediment sites (triangles) such as hydrothermal vents. The relative abundance of Zetaproteobacteria in sediments from other 16S rRNA gene studies was never above 1% and was typically within the range measured from bulk marine sediments from Maine. Sequences are from numerous studies (Supplementary Table S3) that include Sanger, 454, and Illumina sequencing technologies.

Zetaproteobacteria exert a profound impact on global sedimentary biogeochemistry via the production of biogenic, highly-reactive iron oxides despite their low global abundance (∼0.11%)—effectively functioning as keystone organisms in coastal sediments stimulated by burrowing animals. Zetaproteobacteria contribute significantly to the rapid rates of Fe(II) re-oxidation measured and observed in coastal sediments (Chanvalon *et al.*, 2017). Climate change outcomes such as coastal hypoxia may have positive or negative effects on the sedimentary iron biogeochemical cycle—either stimulating microaerobic bacterial iron oxidation resulting in an increase in iron oxide production, thus enhancing dFe release or inhibiting oxidation by the increase in hydrogen sulfide production, precipitating Fe as iron sulfides. The result of an increase or decrease in dFe flux would be enhanced or reduced primary productivity by phytoplankton, respectively. Thus, sedimentary iron oxide formation by Zetaproteobacteria may have a direct impact on important water column processes such as carbon and nitrogen fixation.

## Acknowledgements

This project was funded by a National Science Foundation Biological Oceanography Award number OCE-1459600 (JPB and DE). Sample collection for the Oregon margin and Gulf of Mexico was funded by National Science Foundation grants OCE-1029889 and OCE-1147407 (to JM). We appreciate Sara George for field and laboratory assistance, Peter Larsen for marine polychaete and hemichordate identification, Anton Tramper for sampling worm burrow and sediments from Netherlands, Peter Girguis and David Johnston for helpful discussions, Megan Harder for assistance with iron oxide and DNA extractions, and Matthew Wade for logistical support.

The authors declare no conflict of interest.

---

**Supplementary Figure S1.**
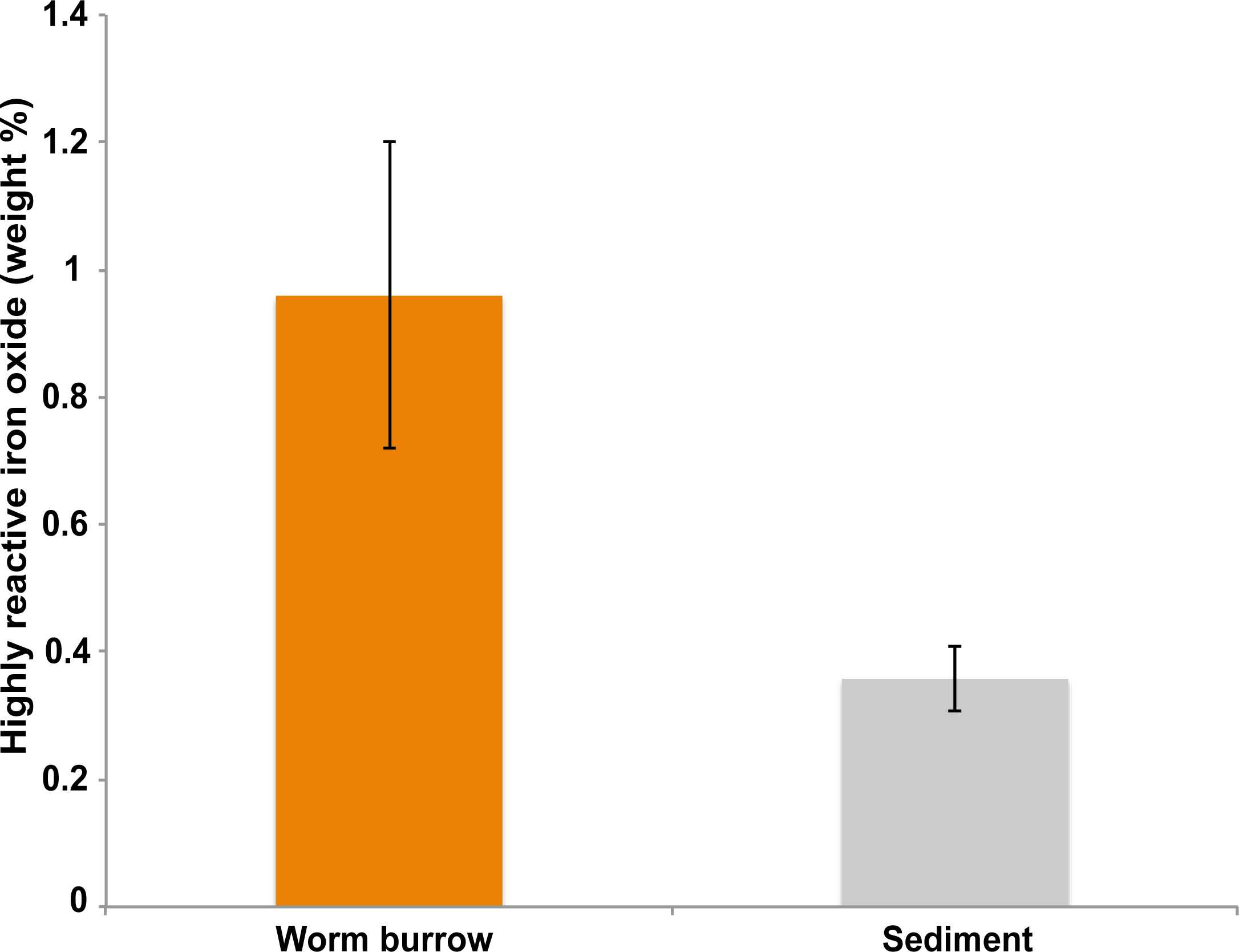
Distribution of highly reactive iron oxides from worm burrows (n=4) and sediments (n=4) at a depth of 3 cm from “The Eddy” Sheepscot River, Maine, USA. Iron oxides are enriched at worm burrow walls as compared to bulk sediments. Error bars represent ± 1 standard deviation from the mean. Highly reactive iron oxides were extracted with sodium dithionite for 2 hours (Poulton and Canfield, 2005).

**Supplementary Figure S2.**
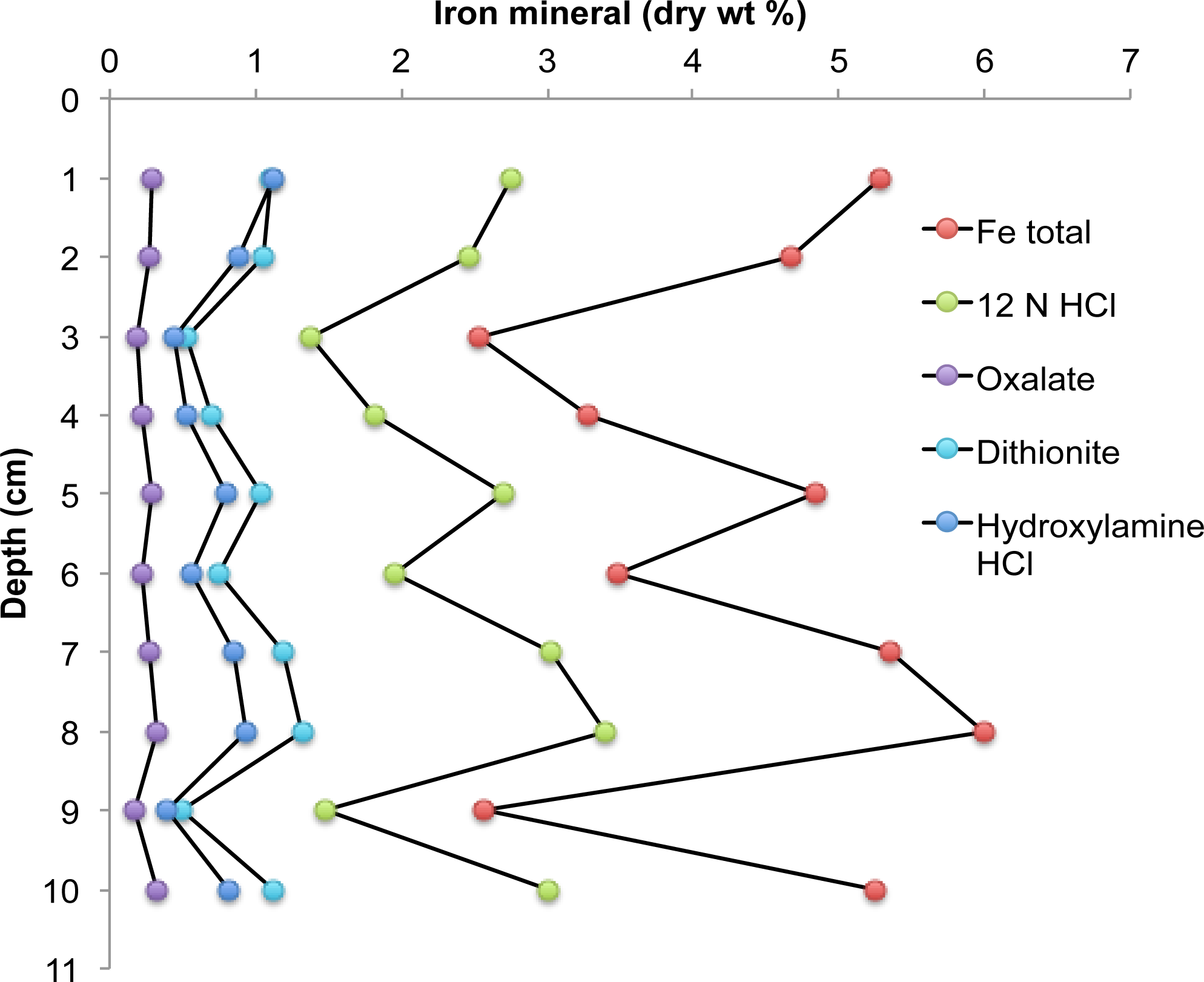
Sequential iron mineral extractions from bioturbated sediments at ‘The Eddy’, Sheepscot River, Maine, USA (sediment temperature= 7 °C; salinity=35 ppt; 43.994827, -69.648632; collected on 1 March 2016). Poorly-crystalline iron oxides—formed primarily by iron-oxidizing Zetaproteobacteria—comprise about 20% of the total iron mineral content in these coastal sediments. Note: dithionite extractions are likely overestimating crystalline iron oxides, as dithionite also liberates Fe from iron sulfides, pyrite, and some sheet silicate iron-containing minerals (Poulton and Canfield, 2005). Hydroxylamine HCl=ferrihydrite and lepidocrocite; dithionite=goethite, hematite, and akaganéite; oxalate=magnetite; 12 N HCl=iron silicates; Fe total=sum of all iron mineral phases.

**Supplementary Figure S3.**
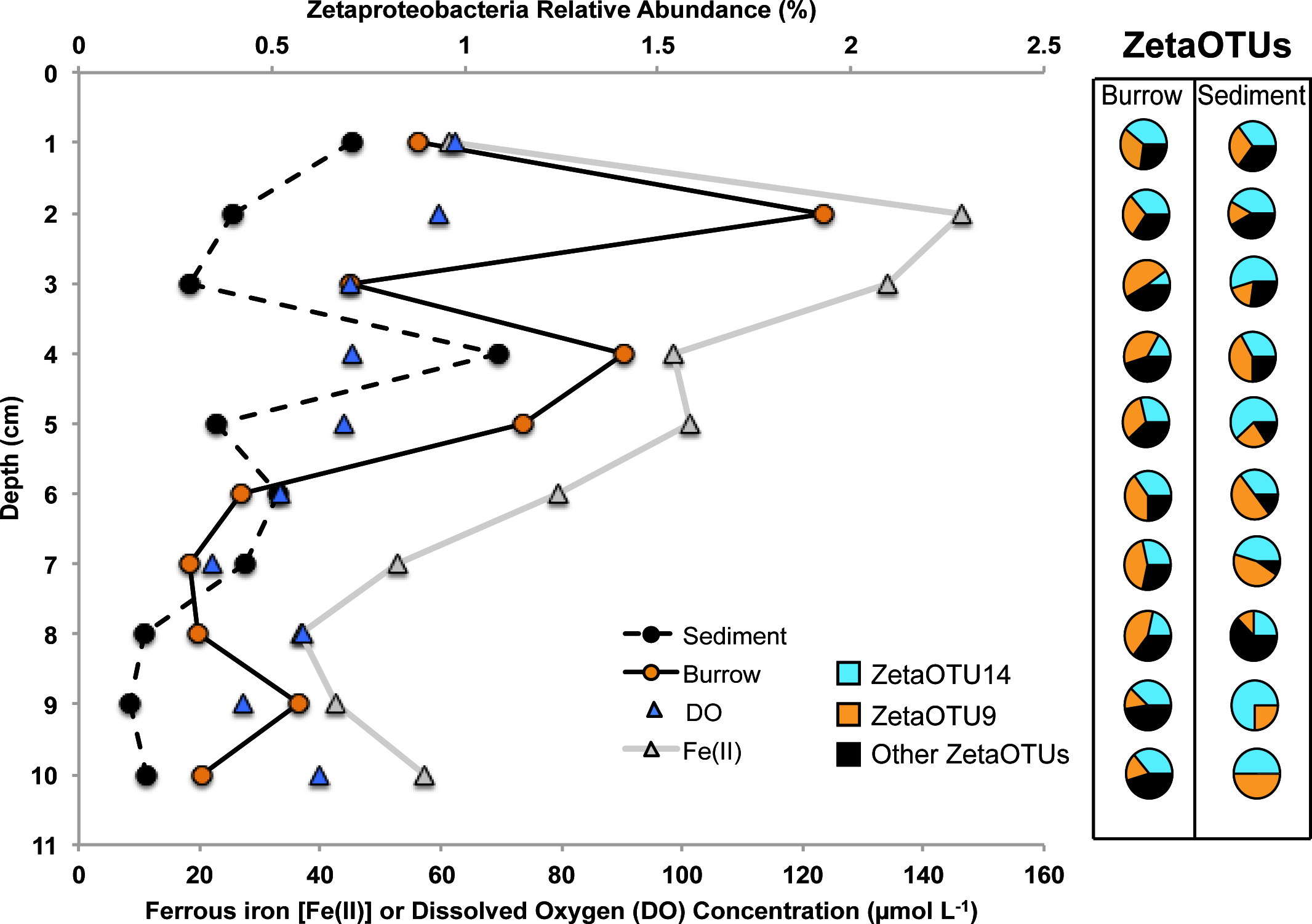
Relative abundance of Zetaproteobacteria in iron oxide-lined burrows (orange cirlces and line) and sediments (black circles and line) as a function of depth from “The Eddy” Sheepscot River, Maine, USA (sampled on 1 March 2016). Pore water ferrous iron concentration (grey circles and line, secondary x-axis) and dissolved oxygen (blue triangles, secondary x-axis) as a function of depth is included for geochemical context. Zetaproteobacteria Operational Taxonomic Units (ZetaOTUs) are indicated on the right from burrows and sediments highlighting dominant ZetaOTUs 14 (blue) and 9 (orange).

**Supplementary Figure S4.**
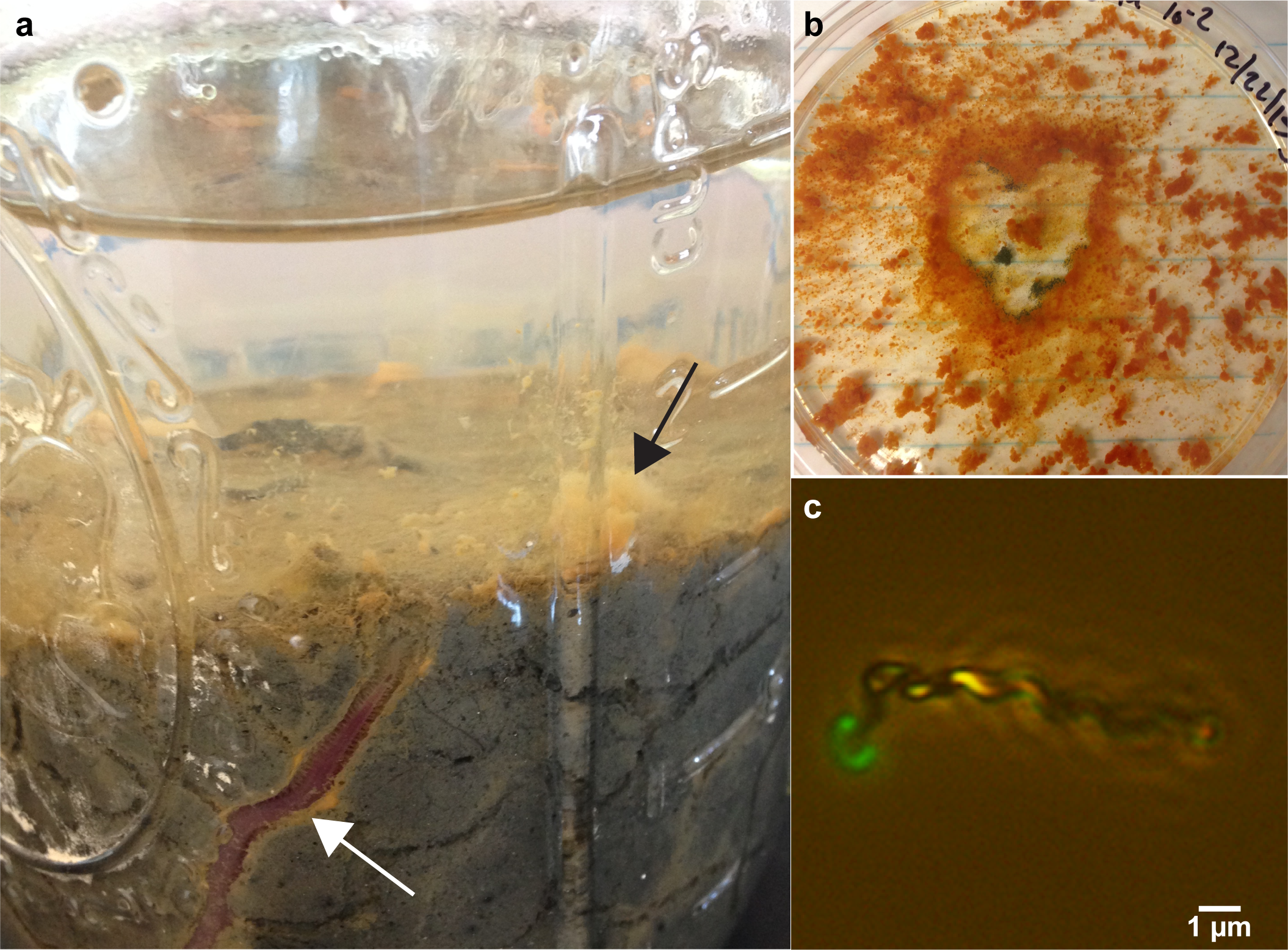
Laboratory bioturbation microcosm (**a**) from a mud flat (Merrow Island) from the Maine coast with abundant iron flocculent (black arrow) forming at the surface of *Glycera dibranchiata* (white arrow) burrows. Zetaproteobacterium strain CSS-1 from ZetaOTU14 growing on zero valent iron (**b**) under low oxygen conditions produces abundant clumps of stalks. Microscope image (**c**) of strain CSS-1 forming an iron oxide stalk stained with Syto13 showing a bean-shaped morphology.

**Supplementary Table S2.** Representative Zetaproteobacteria operational taxonomic units (ZetaOTUs) clustered at 97% nucleotide identity across all worm burrow and sediment samples, and isolate, genome, and iron oxide structure characteristics associated with these ZetaOTUs.

| **ZetaOTU<sup>1</sup>** | **Abundance (%)<sup>2</sup>** | **Isolates/SAGs (#)<sup>3</sup>** | **Biogenic Fe oxide structure<sup>4</sup>** | **Iron Oxidation Metabolism<sup>5</sup>** |
| --- | --- | --- | --- | --- |
| ZetaOTU14 | 31.7 | CSS-1/**4** | twisted stalks | *cyc<sub>2</sub>* (3/4); *cbb<sub>3</sub>* (4/4); *bd* (3/4) |
| ZetaOTU9 | 21.7 | TAG1, SV108/**5** | none | *cyc<sub>2</sub>* (6/7); *cbb<sub>3</sub>* (4/7); *bd* (3/7) |
| ZetaOTU23 | 5.4 | GSB2/**0** | twisted stalks | n.d. |
| ZetaOTU1 | 5.2 | none/**5** | ? | *cyc<sub>2</sub>* (0/5); *cbb<sub>3</sub>* (2/5); *bd* (3/5) |
| ZetaOTU4 | 5.2 | none/**2** | ? | *cyc<sub>2</sub>* (0/2); *cbb<sub>3</sub>* (0/2); *bd* (0/2) |
| ZetaOTU22 | 5.1 | none | ? | n.d. |
| ZetaOTU31 | 2.6 | none | ? | n.d. |
| ZetaOTU11 | 1.5 | JV1, PV1, M34/**0** | twisted stalks | *cyc<sub>2</sub>* (3/3); *cbb<sub>3</sub>* (3/3); *bd* (3/3) |
| ZetaOTU18 | 1.2 | DIS1, CP-8, *M. micogutta*/**0** | twisted stalks/dreads<sup>6</sup> | *cyc<sub>2</sub>* (3/3); *cbb<sub>3</sub>* (3/3); *bd* (3/3) |

<sup>1</sup>ZetaOTU classification using ZetaHunter (github.com/mooreryan/ZetaHunter)
<sup>2</sup>Abundance out of total ZetaOTUs
<sup>3</sup>Single-amplified genomes from Lō'ihi and Mid-Atlantic Ridge deep-sea vent sites (Field et al., 2015; Scott et al., 2015; 2017)
<sup>4</sup>?=no structural information available
<sup>5</sup>*cyc<sub>2</sub>*=putative iron oxidase, *cbb<sub>3</sub>*=high affinity C-type heme copper oxygen reductase, *bd*=high-affinity bd-ubiquinol oxygen reductase, n.d.=no data; number in parentheses indicates genes identified out of total isolate genomes/SAGs.
<sup>6</sup>Field et al., 2016

